# Mechanosensitive Piezo Channels Contribute to Airway Changes in Chronic Obstructive Pulmonary Disease

**DOI:** 10.64898/2026.06.14.732150

**Authors:** Nataliya Migulina, Ben Roos, Theo Borghuis, Maunick Lefin Koloko Ngassie, Li Drake, Wim Timens, Elizabeth R Vogel, Christina M. Pabelick, Corry-Anke Brandsma, Janette K Burgess, Y.S. Prakash

## Abstract

As an intrinsically mechanosensitive organ, the lung experiences a range of mechanical forces. Chronic obstructive pulmonary disease (COPD) involves abnormal macroscopic cellular and extracellular matrix (ECM) changes that impact mechanical properties of the lung. Mechanosensitive Piezo1/2 channels are expressed in the lung including on airway smooth muscle cells (ASM) that mediate cellular responses to stretch and ECM biomechanics. The expression and roles of Piezos in COPD lung ASM are not known. We hypothesized that Piezo expression and activation are altered in COPD lung ASM influencing ECM regulation. Distribution of Piezo proteins in ASM and epithelium of small airways of COPD stage II and IV vs. non-COPD controls was assessed using immunohistochemistry and ImageJ (n=10-17/group). Isolated ASM cells from control (n=6) vs. COPD stage II and IV patients (n=3 each stage) were exposed to stretch or the Piezo1 agonist Yoda1 followed by measurement of ECM gene and protein expression. Less Piezo2 staining was observed in COPD IV patients compared to controls, with lesser area and intensity of staining in the epithelial layer, and lower intensity of staining in ASM and small airways as a whole. Fura-2-based imaging of ASM Ca^2+^ showed lower influx after Yoda1 exposure in COPD II compared to control and COPD IV. Gene expression of Piezo1 increased upon stretching in controls but not in COPD ASM, while Piezo2 protein expression decreased with stretching in all groups. Yoda1 treatment resulted in decreased *collagen1*, *fibulin1* and *periostin* gene and collagen 1 and periostin protein expression in ASM. Overall, these results support a role for Piezo activation in abnormal ECM-ASM cell crosstalk in COPD.

## Introduction

According to the World Health Organisation, chronic obstructive pulmonary disease (COPD) is the third leading cause of death worldwide, causing 23 million deaths in 2019.^1^ Major pathophysiological characteristics that are associated with the decline of lung function and airway obstruction in COPD are inflammation, airway wall remodeling, mucus hypersecretion, and an abnormal increase in air spaces (emphysema).^2^ Importantly, COPD is characterized by changes in extracellular matrix (ECM) and biomechanical properties of the airways and parenchymal tissues^3^ resulting from altered expression of major ECM components such as lower elastin levels in both airways and parenchyma^4,5^, variable changes in collagen expression^4,6,7^ and altered collagen organization^8^. Stabilization and organization of collagen fibers is regulated, in part, by the family of lysyl oxidases (LO; LOX, LOXL1-4). We recently demonstrated higher LOXL1 and lower LOX levels in small airways of COPD patients^9^ suggesting altered biomechanical properties in the context of disease. However, the mechanisms underlying changes in ECM and biomechanical properties of the COPD lung are still under investigation.

Airway remodeling involves thickening and increased fibrosis of the airway smooth muscle (ASM) layer.^10^ ASM cells are involved in contractility as well as production of ECM, inflammatory cytokines, proteases and growth factors^11^ making them a key cell type in airway structure and function. Altered ASM function has been identified in COPD.^12,13^ ASM of COPD patients exposed to cigarette smoke, a major cause of COPD, show greater matrix metalloproteinase 1 and collagen VIII alpha 1 compared to non-COPD ASM.^13^ Conversely, ECM regulates ASM survival, proliferation, migration, contraction and ECM production itself.^14^ Thus, disrupted cross-talk between ASM and ECM can be present in COPD and underlie altered ECM homeostasis and lung tissue remodeling. Therefore, mechanisms underlying such cross-talk represent a novel approach to blunting remodeling.

Several receptors and channels, such as integrins and transient receptor potential (TRP) channels are thought to be involved in the interplay between ECM and ASM.^14,15^ Piezos are a family of mechanosensitive Ca^2+^ permeable ion channels that convert mechanical signals into electrical or chemical signals.^16^ There are two isoforms of which Piezo1 is thought to be more broadly expressed while Piezo2 is known to be expressed in neurons^17^ however the differential expression and functions of Piezo channels are still being investigated. Recent studies have shown that Piezos play important roles in the mechanoregulation of a wide range of physiological and pathological functions across organs.^18^ Piezo channels are expressed in numerous tissues such as lung, bladder, skin and neurons, and play an important role in the regulation of cellular responses through interactions with tissue microenvironmental biomechanical properties and/or ECM components.^19^ For example, increased stiffness of fibronectin-conjugated polyacrylamide substrates promotes activation of dendritic cells through Piezo channels.^20^ In macrophages, Piezos were involved in polarization and actin regulation in response to stiffness^21^, while Piezo1 inhibition in stiffened vascular smooth muscle cells prevents matrix degradation.^22^ In human umbilical cord, mesenchymal stem cell expression of Piezo1 increases with matrix stiffness.^23^ Recent studies have demonstrated that in the absence of ECM proteins, Piezo1 is not sensitive to mechanical forces.^24^ Overall, these data point to mechanosensitive Piezo channels as being important players in airway remodeling in the context of lung disease.

In the lung, Piezo1 is expressed at least in pulmonary endothelial cells^25^, ASM^26^ and alveolar epithelial cells.^27^ In rats, ASM activation of Piezo1 with the agonist Yoda1 decreases cell stiffness and traction force, leading to disruption of stress fibers and impaired horizontal but enhanced vertical cell migration.^26^ Piezo2 has been found in various types of neurons to be important for tactile responses, proprioception, and mechanical pain perception in the lung.^23,28,29^ However, Piezo2 has also been localized to bronchial epithelial cells, macrophages, and smooth muscle cells.^30^ While the lung does express Piezo1 and potentially Piezo2, its role in airway remodeling in COPD has not been investigated. In this study, we investigated expression patterns of Piezo channels in small airways in COPD, with a focus on ASM. We explored the effect of stretch on ASM Piezo channel expression and the impact of Piezo activation including expression of ECM and ECM-related genes and proteins.

## Materials and Methods

### Human lung samples

Human lung tissues were obtained from patients undergoing surgery for resection of focal lung tumors or lung transplantation for severe COPD at University Medical Center Groningen (UMCG). Lung function was assessed prior to surgery, and patients were classified as non-COPD when forced expiratory volume in one second (FEV1)/forced vital capacity (FVC) was > 70% and FEV1 ≥ 80%, GOLD stage II as FEV1/FVC < 70% and 50% < FEV1 < 80% and GOLD stage IV as FEV1/FVC < 70% and 30% < FEV1 < 50% predicted. In non-transplant samples, to avoid possible disease effects, lung tissues were sampled as distant from the tumor as possible. The study protocol was consistent with the Research Code of UMCG (umcgresearch.org) and national ethical and professional guidelines (“Code of conduct for Health Research: Gedragscode-Gezondheidsonderzoek-2022.pdf (coreon.org)). Lung tissues were derived from leftover lung material after lung surgery from archival materials that are exempt from consent in compliance with applicable laws and regulations (Dutch laws: Medical Treatment Agreement Act (WGBO) art 458 / GDPR art 9/ UAVG art 24). This material was not subject to the Medical Research Human Subjects Act in the Netherlands, as was confirmed by a statement from the Medical Ethical Committee of the UMCG. All donor material and clinical information were deidentified prior to experimental procedures, blinding any identifiable information to the investigators.

### Histology and immunohistochemistry

Deparaffinized lung tissue sections were incubated with citrate buffer (10 mM sodium citrate, pH 6) for 15 min at 100°C for antigen retrieval. The slides were cooled for 30 min at room temperature (RT) and washed with Phosphate Buffered Saline (PBS). Then, the slides were incubated with monoclonal mouse anti-human Piezo1 antibody (1:200, Novus Biologicals, NBP2-75617, Cambridge, UK) or polyclonal rabbit anti-human Piezo2 antibody (1:100, Novus Biologicals, NBP1-78624, Cambridge, UK) for 1.5h followed by anti-mouse or anti-rabbit horseradish peroxidase-conjugated secondary antibody (1:100, DAKO, Glostrup, Denmark) at RT for 30 min. NovaRED (Vector Laboratories, SK-4800, Burlingame, CA, USA) was used as the chromogen and hematoxylin as a nuclear counterstain. Finally, tissue sections were dried at RT and mounted usig using a permanent mounting medium.

### Image analyses

Tissue sections were scanned using a Hamamatsu slide scanner (Nonozoomer 2.0 HT, Hamamatsu Photonics, Shizuoka, Japan). Slides were analyzed with ImageJ (FIJI version 2.9.0)^31^ to quantify the density and distribution of staining. Positive staining was identified by the separation of the original images into blue (hematoxylin-image), and red (NovaRED-image) using the color deconvolution plugin by Landini.^32^ To determine the correct optical density vectors of Hematoxylin and NovaRED we followed the protocol previously described by Ruifrok et al.^33, 34^ Macros were used to process the images and to generate automated numbers of pixels in the different images. The image analyses calculated the average intensity of the positive pixels above the threshold (mean staining intensity - formula 1) and the percentage of positively stained tissue area (Area % - formula 2).

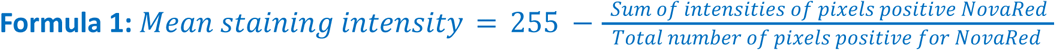

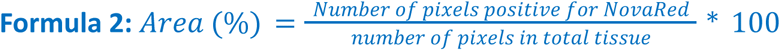

From 17 non-COPD, 10 COPD II and 10 COPD IV patients, tissue sections were stained separately for Piezo1 and Piezo2. Up to six airways in each tissue section were selected for analyses (Figure 1A) (two to six small airways per section). Within each airway, three regions of interest were manually selected using Photoshop and then analyzed: the small airway as a whole (Figure 1B), the epithelial layer (Figure 1C) and the ASM (Figure 1D) within the airway wall.

**Figure 1.**
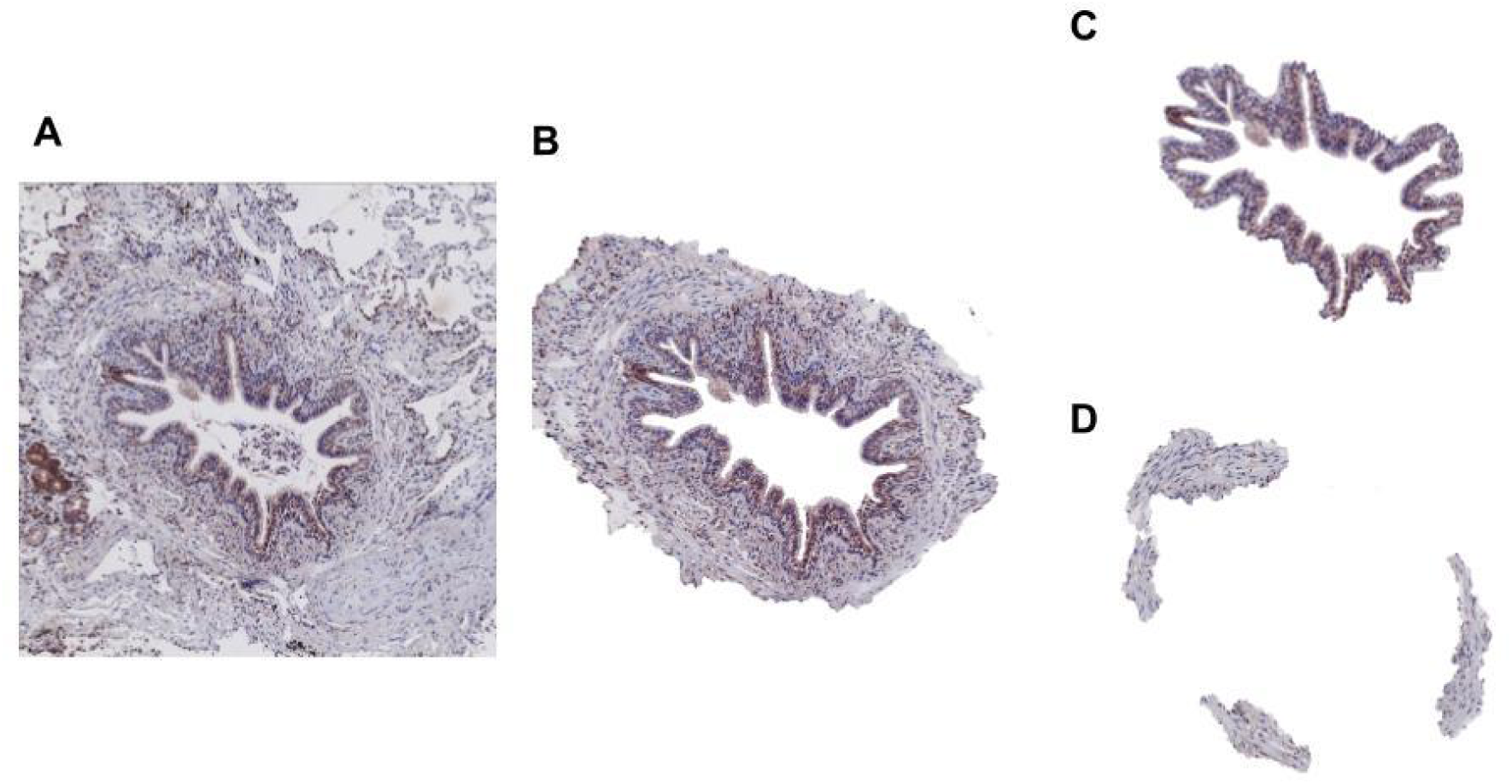
Preparation of images for analysis of specific structures in the human airways. Representative images of immunohistochemically stained human airway tissues from a COPD stage IV patient (A) and (red = protein of interest; blue = cytoplasm and nuclei). The regions isolated for analyses were total airway (B),the airway epithelial layer (C) and airway smooth muscle (D).

### Airway smooth muscle cell culture

Human ASM cells were isolated as previously described.^35, 36^ 3^rd^–6^th^ level bronchi were obtained from lung samples of patients undergoing thoracic surgery at Mayo Clinic Rochester, MN for focal, non-infectious tumors (all protocols approved by Mayo Clinic Institutional Review Board, IRB# 16-09655). Patient consents (written or video/verbal) were obtained during clinic visits prior to surgical decisions, and relevant clinical data collected upon acquisition of lung tissues. For anonymization/de-identification and storage, samples were assigned unique numbers unrelated to any patient identifier (patient number, age, sex, date of birth, date of surgery, etc.). COPD patients were classified as follows according to the severity of airflow limitation (1) non-COPD; FEV1/FVC ≥ 70% and FEV1 ≥ 80%; (2) COPD; FEV1/FVC < 70%. Patients with asthma, infectious diseases, or interstitial lung diseases were not included. Following tissue collection, bronchi were dissected, the epithelial layer removed using tweezers and the ASM bundles dissected, immersed in ice-cold Hanks’ Balanced Salt Solution (HBSS; 2 mM Ca^2+^) and cells enzymatically dissociated. All cells were used between passages 3 and 4. For all cell experiments, except for protein ECM expression, ASM cells were obtained from 12 patients; 6 non-COPD control patients and 6 patients with COPD (3 COPD stage II and 3 COPD stage IV) (group 1). For ECM protein expression, a different set of ASM cells was used, obtained from 10 patients: 5 non-COPD control patients and 5 patients with COPD (4 COPD stage II and 1 COPD stage IV). Patients’ demographics are in Table 1.

**Table 1.**
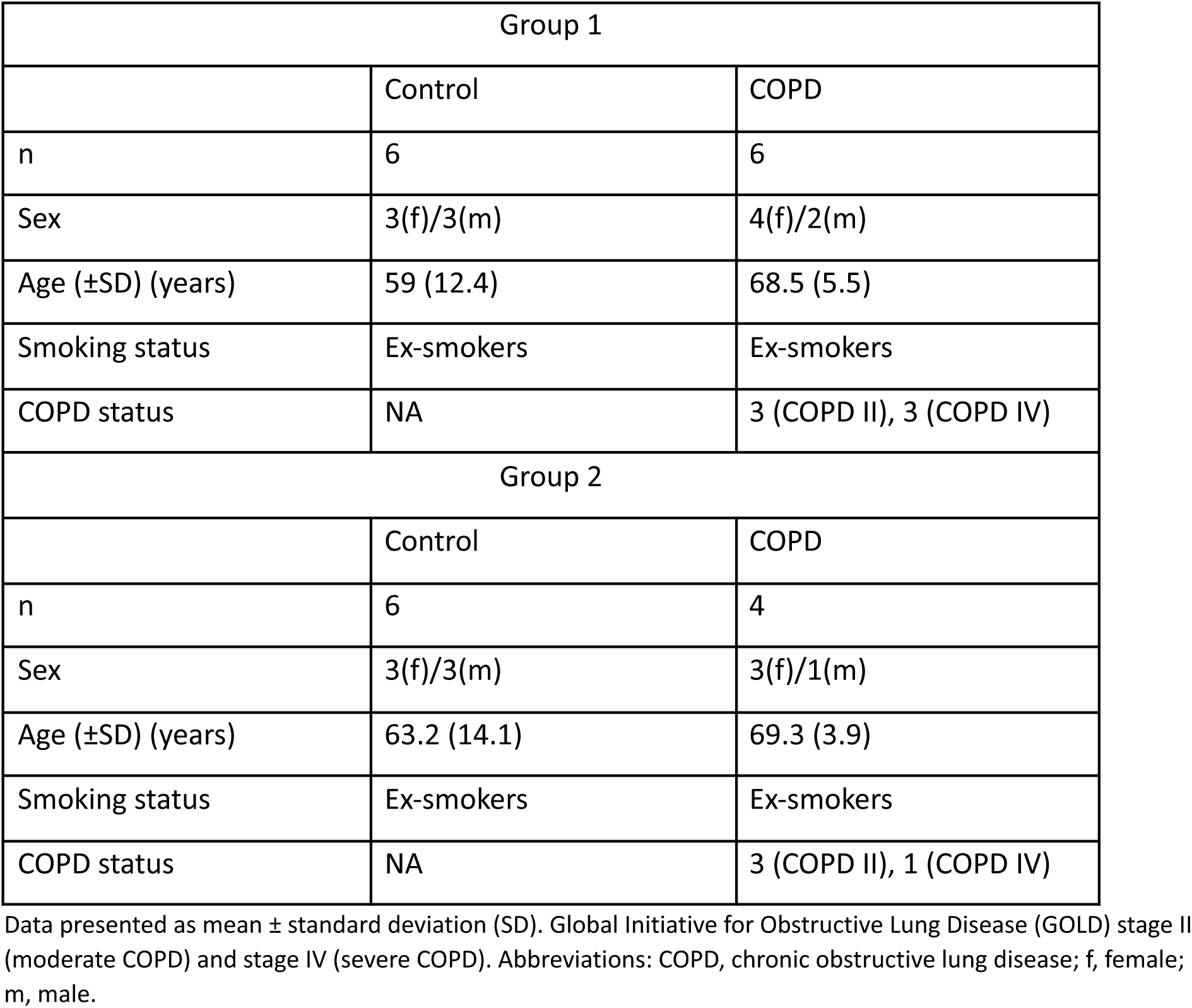
Patient demographics for ASM.

| Group 1 |  |  |
| --- | --- | --- |
|  | Control | COPD |
| n | 6 | 6 |
| Sex | 3(f)/3(m) | 4(f)/2(m) |
| Age (±SD) (years) | 59 (12.4) | 68.5 (5.5) |
| Smoking status | Ex-smokers | Ex-smokers |
| COPD status | NA | 3 (COPD II), 3 (COPD IV) |
| Group 2 |  |  |
|  | Control | COPD |
| n | 6 | 4 |
| Sex | 3(f)/3(m) | 3(f)/1(m) |
| Age (±SD) (years) | 63.2 (14.1) | 69.3 (3.9) |
| Smoking status | Ex-smokers | Ex-smokers |
| COPD status | NA | 3 (COPD II), 1 (COPD IV) |

Data presented as mean ± standard deviation (SD). Global Initiative for Obstructive Lung Disease (GOLD) stage II (moderate COPD) and stage IV (severe COPD). Abbreviations: COPD, chronic obstructive lung disease; f, female; m, male.

### ASM Piezo activation and stretch

Cells for ECM analysis were grown in F12 DMEM with L-glutamine, 10% fetal bovine serum (FBS; Sigma-Aldrich, St. Louis, MO, USA), 1% antibiotics and antimycotic (AA; Gibco, lot no. 2441424)and incubated at 37°C, 5% CO2. After cells reached confluence, ASM were trypsinized and seeded in 6-well culture plates (Corning Inc., Corning, NY, USA) at a density of 10,400 cells/cm^2^. After 24 h media was replaced with F12 DMEM with low serum (0.5% FBS).

To investigate the functional activity of Piezo1 in regulating ECM production, 24 h after seeding ASM cells were stimulated with 1 µM Yoda1 (2-[5-[[(2,6-Dichlorophenyl)methyl]thio]-1,3,4-thiadiazol-2-yl]-pyrazine; Biotechne, Minneapolis, MN, USA, Cat. 5586) a selective chemical Piezo1 activator^37^ followed by RNA and protein collection after 24 h and 48 h, respectively to measure gene and protein expression of ECM elements.

To determine the effect of mechanical forces on Piezo expression, ASM were placed in an *in vitro* cell stretcher system and stretched for 24 h and 48 h for RNA and protein collection respectively. Cells were seeded in 6-well plates with flexible membranes (BioFlex, Culture Plates, FlexCell, Burlington, NC, USA) at a concentration of 6,250 cells/cm^2^. First, cells were placed in the middle of a well in 500 µL growth medium so the medium with cells did not touch the walls of the well and allowed to attach. After 2 h, 1.5 mL of growth medium was added and plates were placed in a cell stretcher system (FX-6000T Tension System, FlexCell, Burlington, NC, USA) such that the membranes were stretched over a rigid post following negative pressure applied to the sealed unit in which the plates were placed. Cells underwent an oscillatory stretch of 15% at 0.166 Hz (mimicking breathing). 24 h after commencing stretching, samples for RNA analyses were collected in RLT lysis buffer (Qiagen, Hilden, Germany) containing 10 μL of beta-mercaptoethanol (Gibco, Carlsbad, CA, USA) per 1 ml. Proteins were collected 48 h after initiating oscillatory stretch. Once media was removed, cells were placed on ice and washed twice with cold PBS and cold RIPA Buffer (Cat. No. 89901, Thermo Fisher) containing Protease Inhibitor Cocktail (Cat. No. 78410, Thermo Fisher) and Halt™ Phosphatase Inhibitor Cocktail (Cat. No. 78420, Thermo Fisher). The lysate was collected using a cell scraper and transferred to a microcentrifuge tube before being sonicated for 20 seconds and centrifuged at ∼14,000 × g for 15 minutes. Cell supernatant was collected and stored at -80 °C. Cells in control non-stretched plates underwent the same procedure but instead of a cell stretcher system, they were grown in a regular incubator at 37°C and 5% CO_2_.

### Gene expression analysis

RNA was purified using a RNeasy Mini Kit (Cat. No.74004, Qiagen, Hilden, Germany) according to the manufacturer’s protocol. The concentration and quality of RNA were assessed by using NanoDrop Spectrophotometer (Thermo Fisher, Waltham, MA, USA). Collected RNA was stored at -80°C. cDNA was synthesized from RNA using the Transcriptor First Strand cDNA Synthesis Kit (Roche, Indianapolis, IN, USA) according to the manufacturer’s instructions. A solution containing 100 ng RNA sample in a total volume of 20 µL per sample was placed in ThermoCycler (Bio-Rad, Hercules, CA, USA) for denaturation for 10 min at 65°C, then for cDNA synthesis 10 min at 25°C followed by 30 min at 55°C and 5 min at 85°C. After running the protocol, cDNAs were stored at −20°C. SYBR Green PCR Master Mix (Roche; 4707516001) was used for qPCR. The PCR program sequence included two steps. For step one samples were placed for 60 sec at 95°C; step two included 40 cycles with the sequence of 15 sec at 95°C, 15 sec at 60°C and 15 sec at 72°C. Data were analyzed using the comparative ΔΔCt method and compared to *16S* as the housekeeping gene. Predesigned human primers (Integrated DNA Technologies, Coralville, IA, USA) that were used are listed in Table 2.

**Table 2.** List of predesigned primers.

| Gene name | Assay ID |
| --- | --- |
| 16S | Hs.PT.58.366374 |
| LOX (*Lysyl Oxidase*) | Hs.PT.58.1471308 |
| *LOXL1 (Lysyl Oxidase like 1)* | Hs.PT.58.40240878 |
| *LOXL2 (Lysyl Oxidase like 2)* | Hs.PT.58.2256221 |
| *Col1A1 (Collagen1)* | Hs.PT.58.15517795 |
| *Col3A1 (Collagen3)* | Hs.PT.58.4249241 |
| *BMP1 (Bone Morphogenetic Protein 1)* | Hs.PT.56a.25570664 |
| *Periostin* | Hs.PT.58.4452022 |
| *Fbln1 (Fibulin 1)* | Hs.PT.58.45318636 |
| *Piezo1* | Hs.PT.58.14351120 |
| *Piezo2* | Hs.PT.58.24452135 |

### Protein measurement

Protein quantification was performed using BCA protein assays (Thermo Fisher #89901). Protein expression levels were measured using JESS, a capillary-based electrophoresis system (Protein Simple, San Jose CA, United States). Following the manufacturer’s instructions, 0.3 µg protein was loaded into 12-230-kDa or 66-440-kDa JESS separation modules with appropriate primary and secondary antibodies validated for that system. The primary antibodies used were Piezo1 (LSBio, LS-C751636, 1:50 dilution), Piezo2 (Novus Biologicals, NBP2-61130, 1:25 dilution), Periostin (Origene, CF804575, 1:50 dilution), Collagen 1 (Col1, Novus, NBP1-30054, 1:50 dilution), Collagen 3 (Col3, Abcam, 7778, 1:50 dilution). Bone morphogenetic protein 1 (BMP1, abcam, ab205394, 1:50 dilution), Vinculin (Cell Signaling Technology, 13901, 1:50 dilution). The secondary antibodies were purchased from ProteinSimple: Anti-Mouse Secondary HRP Antibody (# 042-205), Anti-Goat Secondary HRP Antibody (# 043-522-2), and Anti-Rabbit Secondary HRP Antibody (# 042-206). Total Protein Detection Module for Chemiluminescence based total protein assays (# DM-TP01) were used to determine the total protein expression. Digital representations of the electropherograms were used and then quantified using Compass for Simple Western Software. Expression of Piezo proteins was normalized to cytoskeletal protein vinculin, whereas expression of ECM-related proteins was normalized to total protein.

### Measurement of intracellular Ca^2+^ concentration ([Ca^2+^]_i_)

ASM were seeded in 8-well chambers (Lab-Tek, Rochester, NY) in F12 DMEM containing 10% FBS and 1% P/S at a density of 3750 cells/cm^2^. Cells were grown to 50% confluence and serum starved in F12 DMEM with L-glutamine and 1% P/S for 24 h prior to experimentation. Cells were washed 3 times with freshly prepared HBSS and incubated with Ca^2+^ indicator Fura-2/AM (5 μM; ThermoFisher Scientific, Scotts Valley, CA) for 30 min at room temperature. Cells were then washed with HBSS. Fura-2 loaded ASM were then imaged using a 40 × lens on a Nikon Eclipse Ti-U microscope using HROMA 79000V2 filters (Chroma Technology Corp., Bellows Falls, VT, USA). Cells were perfused with HBSS for 2 min, followed by treatment with or without Yoda1 (1 µM) for 6 minutes and perfused again with HBSS for a further 7 min. During the 15 min of perfusion, 340/380 nm emission ratio of Fura-2/AM was recorded using a Princeton Instruments/Hamamatsu CCD camera. Images from an area containing 8-15 cells per well and 3 wells per patient were recorded.

### Statistical analysis

Intracellular Ca^2+^ concentration analyses were performed using SPSS Statistics software, IBM (version 27). IHC staining data analyses were performed using R (R version 4.5.1). IHC staining results included multiple small airways, epithelial layers and ASM layer measurements for each patient, and therefore were analyzed using a linear mixed effects regression model. The same model was used for Ca^2+^ concentration because measurements included multiple ASM cells per patient. The differences in gene and protein expression between non-COPD and COPD groups were analyzed using the Mann-Whitney U test, and within non-COPD and COPD groups with the Wilcoxon test. All data are expressed as a median with 95% confidence interval. P < 0.05 was considered significant.

## Results

### Piezo proteins are expressed in human lung tissue

IHC was used to examine Piezo expression and localization in human lung tissue samples containing airways (20 patients, non-COPD ex-smokers and ex-smoker patients with severe COPD (GOLD stage IV; n = 10 per group; Table 3).

**Table 3.** Patient demographics table.

|  | Control ex-smokers | COPD stage II/III ex-smokers | COPD stage IV ex-smokers |
| --- | --- | --- | --- |
| N | 17 | 10 | 10 |
| Sex | 8(f)/9(m) | 2(f)/8(m) | 7(f)/3(m) |
| Age (± SD) (years) | 65.8 (10.1) | 74.9 (5.1) | 50.8 (2.9) |
| FEV1/FVC % (± SD) | 76.2 (5.6) | 57.0 (7.6) | 25.1 (6.2) |
| FEV1% predicted (± SD) | 101.4 (18) | 61.4 (7.6) | 19.0 (6.2) |

Data presented as mean ± standard deviation (SD). Abbreviations: BD, bronchodilator; f, female; m, male; FEV1, forced expiratory volume in 1 s; FVC, forced vital capacity; GOLD, Global Initiative for Chronic Obstructive Lung Disease.

Visual examination of staining patterns showed that both Piezo1 and Piezo2 are expressed in bronchial epithelial cells and stronger on their apical region. Both Piezo isoforms are expressed in ASM, vascular smooth muscle cells and endothelial cells (Figure 2 A-D).

**Figure 2.**
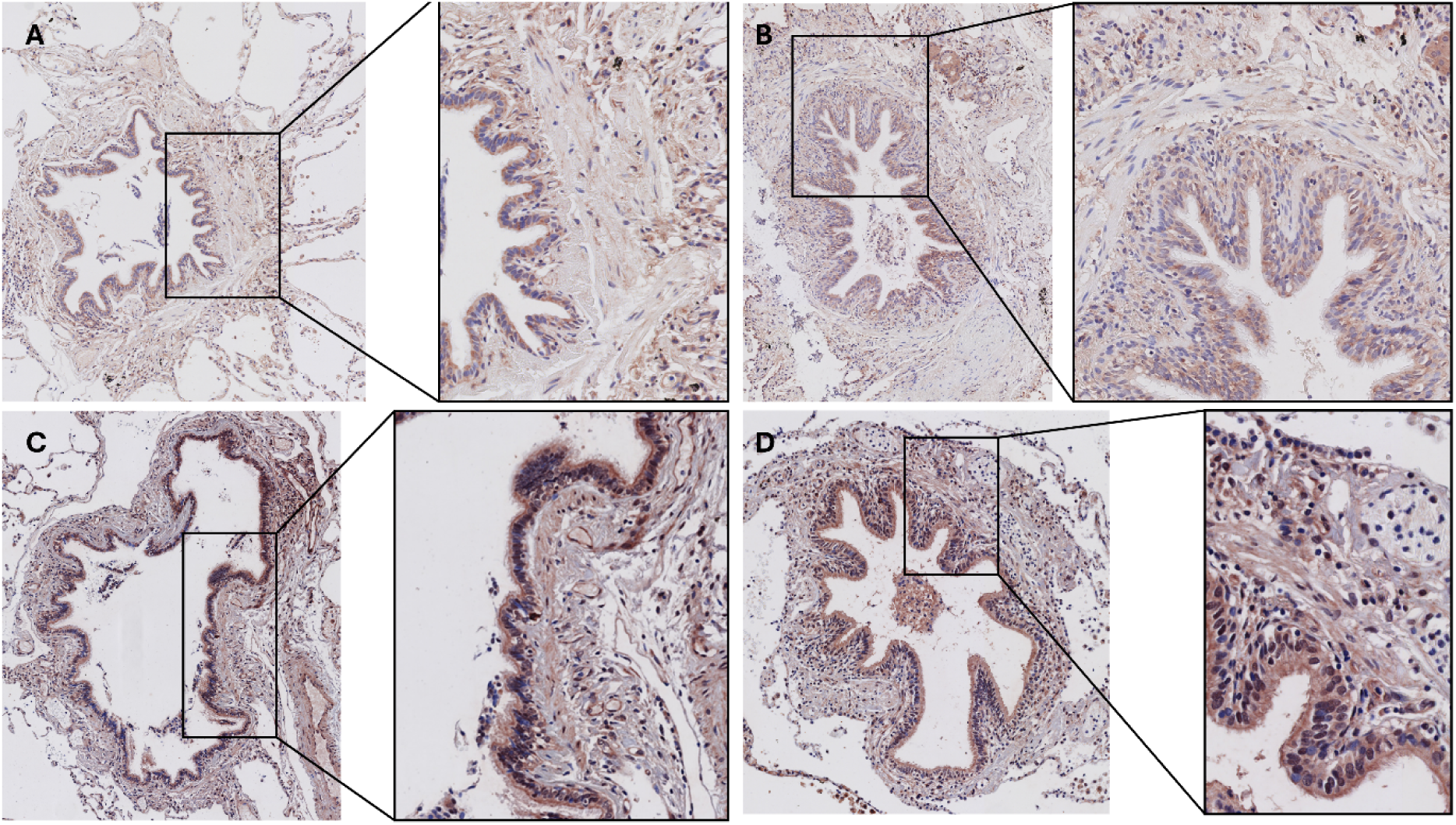
Representative expression of Piezo1 and Piezo2 in lung tissue. Piezo1 in non-COPD (A) and COPD IV lung tissue (B), and Piezo2 in non-COPD (C) and COPD IV tissues (D) were constitutively expressed in the epithelial cells, ASM, and vascular smooth muscle and endothelial cells. The displayed images are representative of the staining results obtained in the other patients, with similar patterns observed. Red/brown = Piezo1 or 2; blue = hematoxylin.

### Lower expression of Piezo2 in COPD small airways

The percentage area of positive pixels and the average staining intensity of Piezo1 showed no differences between groups across the different compartments of the small airways (Figure 3 A-F). Whereas the percentage area of Piezo2 staining in small airways did not differ significantly between groups (Figure 3 G), the staining intensity was lower in the small airways of COPD IV patients compared to the non-COPD group (Figure 3 H). In the epithelial layer of COPD IV patients, both the percentage area and staining intensity of Piezo2 staining was lower compared to the non-COPD group (Figure 3 I, J). While the percentage area of Piezo2 staining in ASM cells showed no significant difference, its staining intensity was lower in COPD IV ASM cells relative to the non-COPD group (Figure 3 K, L). In the COPD II group, Piezo2 expression did not significantly differ from either the non-COPD or COPD IV groups (Figure 3 G-L).

**Figure 3.**
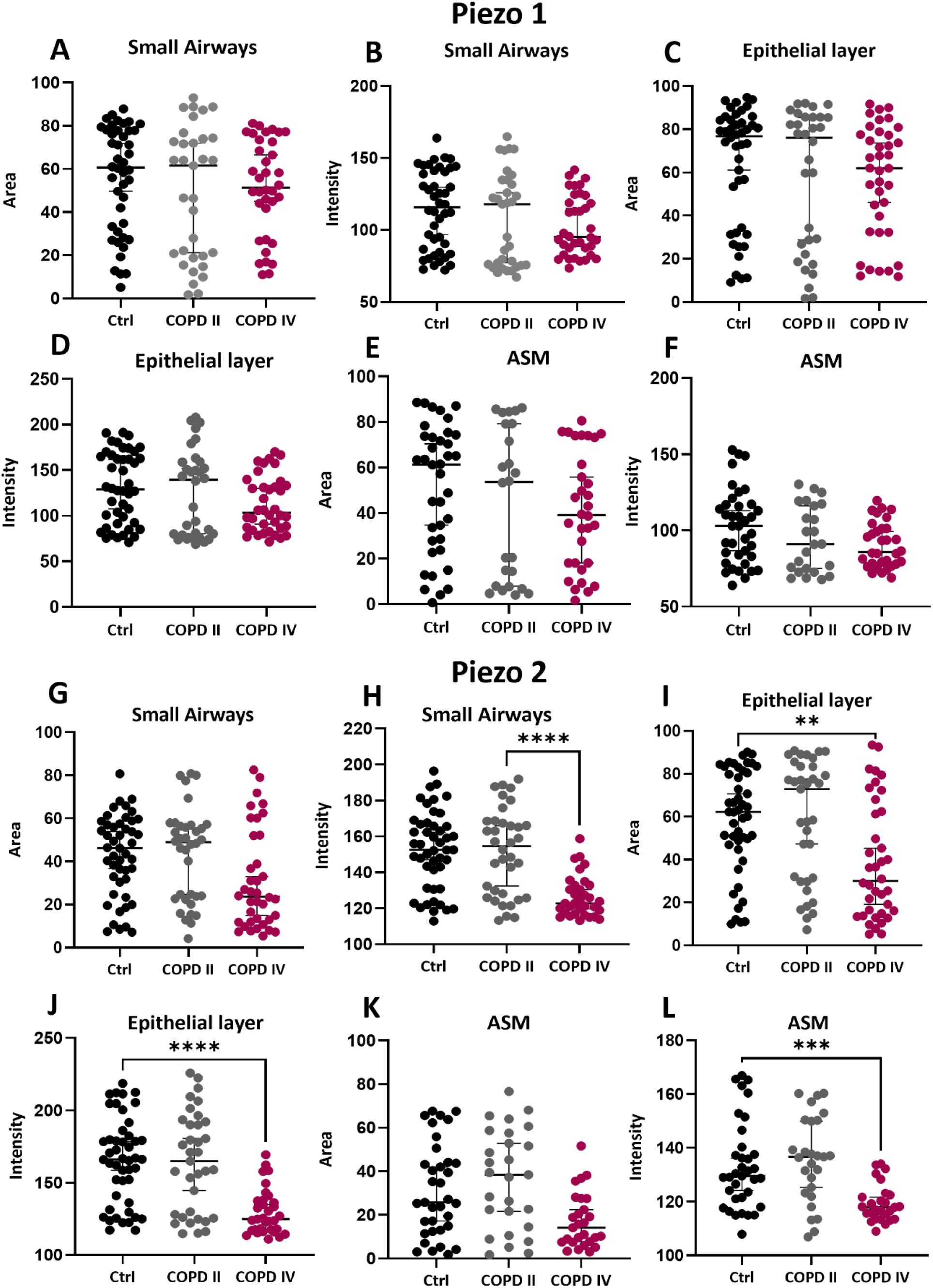
The image analysis of Piezo1 and 2 immunohistochemistry was performed to calculate the percentage area of positive pixels and average staining intensity. Each lung tissue section contained from two to six small airways. Within each airway, three regions of interest were selected: the total small airway, the epithelial layer and the airway smooth muscle (ASM). Piezo1 staining: percentage area of positive pixels in the small airway (A), average intensity of staining in the small airway (B), percentage area of positive pixels in the epithelial layer (C), average intensity of staining in the epithelial layer (D), percentage area of positive pixels in ASM (E), average intensity of staining in ASM (F); Piezo2 staining: percentage area of positive pixels in the small airway (G), average intensity of staining in the small airway (H), percentage area of positive pixels in the epithelial layer (I), average intensity of staining in the epithelial layer (J), percentage area of positive pixels in ASM (K), average intensity of staining in ASM (L). Data are expressed as median with 95% confidence interval. Data analyzed using linear mixed-effects regression model; *p < 0.05.

### Stretch upregulates *Piezo1* gene expression in ASM

To begin to understand if mechanical forces impact the response of Piezo in ASM, we investigated the effect of oscillatory stretch on *Piezo* gene expression in ASM cells from the combined COPD and non-COPD group. Stretched ASM showed higher expression of *Piezo1* compared to non-stretched cells (Figure 4A), but there was no difference in *Piezo2* expression (Figure 4B). Next, we investigated whether *Piezo* gene expression was different between COPD versus non-COPD ASM and whether the effect of stretch differed between groups. There was no difference in *Piezo1* expression between static non-COPD and COPD ASM. Stretched non-COPD cells showed increased *Piezo1* expression compared to their non-stretched control, whereas the effect of stretch was not significant in COPD ASM (Figure 4C). There was no difference in *Piezo2* expression between non-COPD and COPD ASM. Stretching did not change *Piezo2* expression in either the non-COPD or COPD groups (Figure 4D).

**Figure 4.**
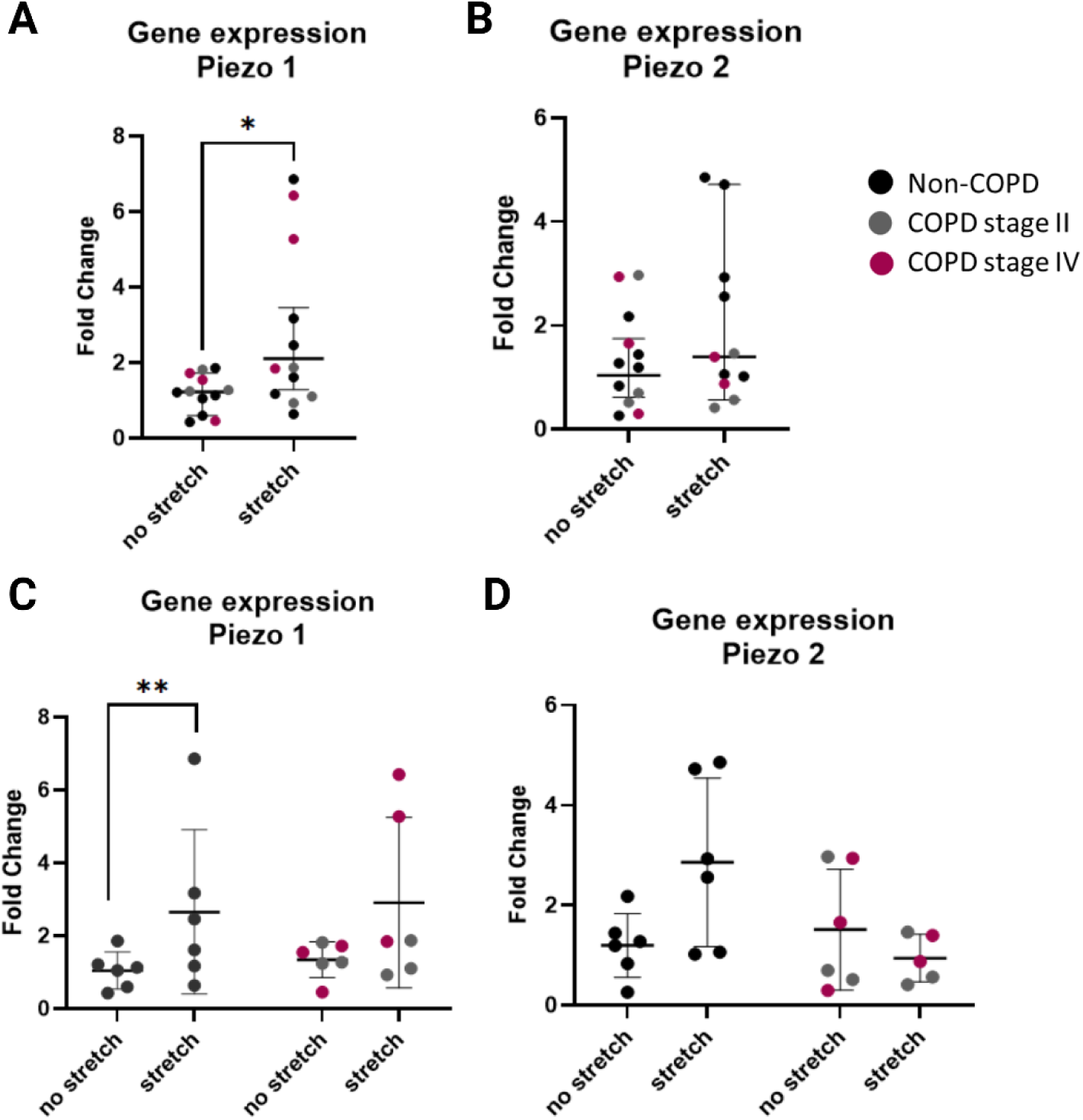
Piezo gene expression in ASM from non-COPD and COPD patients and the influence of stretch. ASM were placed in the cell stretcher system with an oscillatory stretch of 15% at 0.166 Hz for 24 hours. Gene expression was analyzed by using qPCR. Results are presented as gene expression of *Piezo1* and *Piezo2* relative to *16S* of non-COPD and COPD ASM with and without stretching. (A) *Piezo1* expression in ASM with and without stretch (n=12); (B) *Piezo2* expression in ASM with and without stretch (n=12); (C) effect of cellular origin (COPD or non-COPD) on *Piezo1* gene expression in non-stretched and stretched ASM (n=6); (D) effect of cellular origin (COPD or non-COPD) on *Piezo2* gene expression in non-stretched and stretched ASM (n=6). Data are expressed as median with 95% confidence interval. The difference between stretched and non-stretched groups in ASM was analyzed using the Wilcoxon test; the difference between non-COPD and COPD ASM was analyzed using the Mann-Whitney test. *p < 0.05.

### Stretch downregulates Piezo2 protein expression in ASM

Initially, we investigated the effect of oscillatory stretch on ASM Piezo protein expression in the combined COPD and non-COPD group. Stretch did not change Piezo1 protein levels compared to non-stretched cells (Figure 5A). However, levels of Piezo2 protein were downregulated after stretch (Figure 5B). When considering the differences between COPD and non-COPD ASM cells, we found no differences in Piezo1 or Piezo2 protein levels between non-COPD and COPD ASM under static or stretched conditions (Figure S1 A, B).

**Figure 5.**
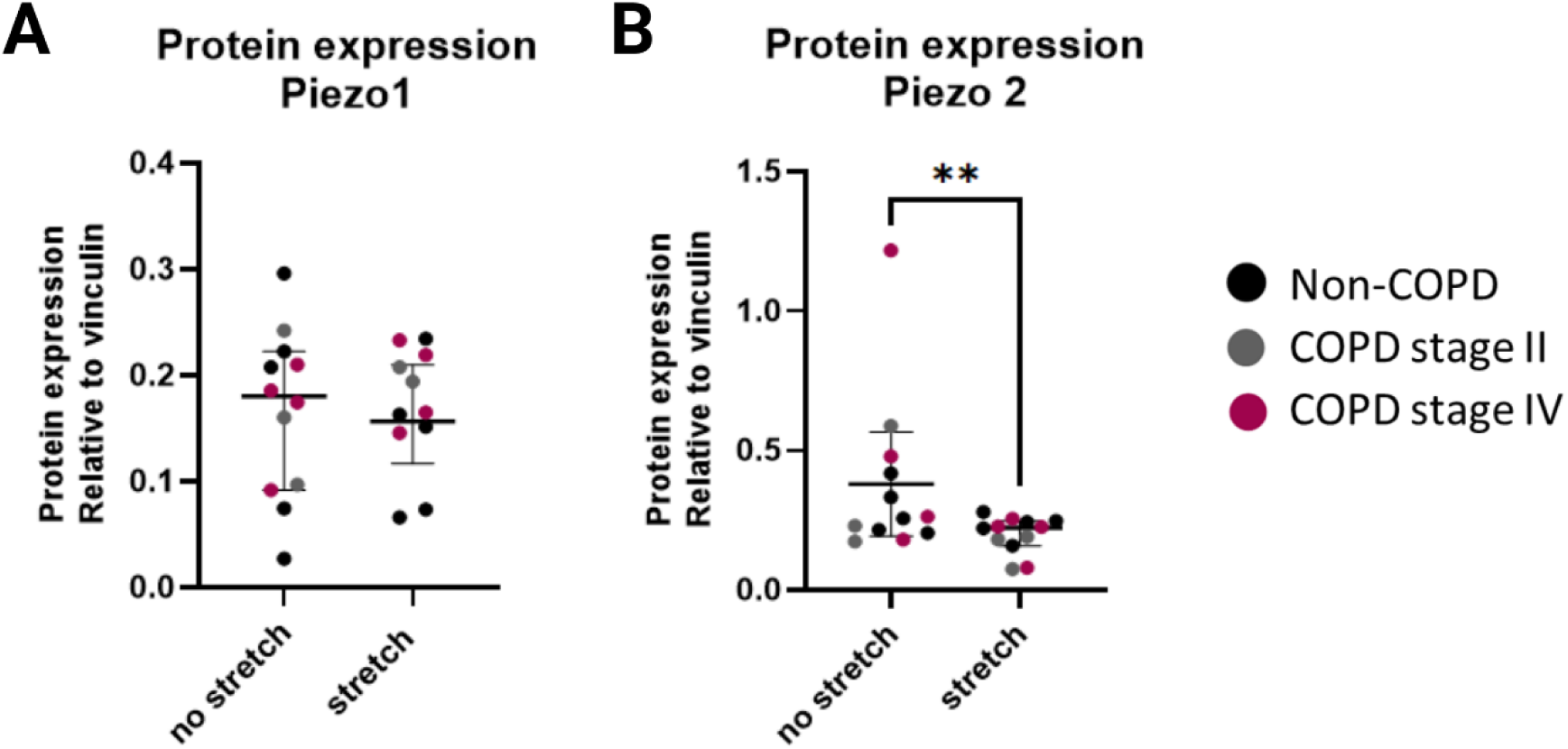
The effect of stretch on Piezo protein levels in ASM cells. ASM were stretched in a cell system FlexCell for 48 hours. Protein expression of Piezo in non-stretched and stretched ASM was analyzed using the WES system. Piezo expression was calculated relative to levels of vinculin protein. (A) Piezo1 protein expression in ASM derived from both non-COPD and COPD patients (n=12); (B) Piezo2 protein expression in ASM derived from both non-COPD and COPD patients (n=12). Data are expressed as median with 95% confidence interval. The difference between non-stretched and stretched groups was analyzed using the Wilcoxon test. *p < 0.05.

### Activation of ASM Piezo1 induces lower Ca^2+^ responses in COPD II

Yoda1 activates Piezo1 in the absence of mechanical stimuli. To investigate the functional activity of the Piezo1 channel in COPD compared to non-COPD ASM, cells from non-COPD and COPD II/COPD IV patients were stimulated with Yoda1 and changes in [Ca^2+^]_i_ were recorded (Figure 6 A, B). Ca^2+^ influx upon Yoda1 stimulation varied between patients and we observed that ASM cells derived from COPD II patients are less responsive to Yoda1 stimulation compared to COPD IV and non-COPD patients. The amplitude of the Ca^2+^ response to Yoda1 was calculated as a difference between the peak value and the baseline value prior to Yoda1 addition. The response amplitude of COPD II ASM cells treated with Yoda1 was significantly lower compared to COPD IV or non-COPD ASM (Figure 6 C).

**Figure 6.**
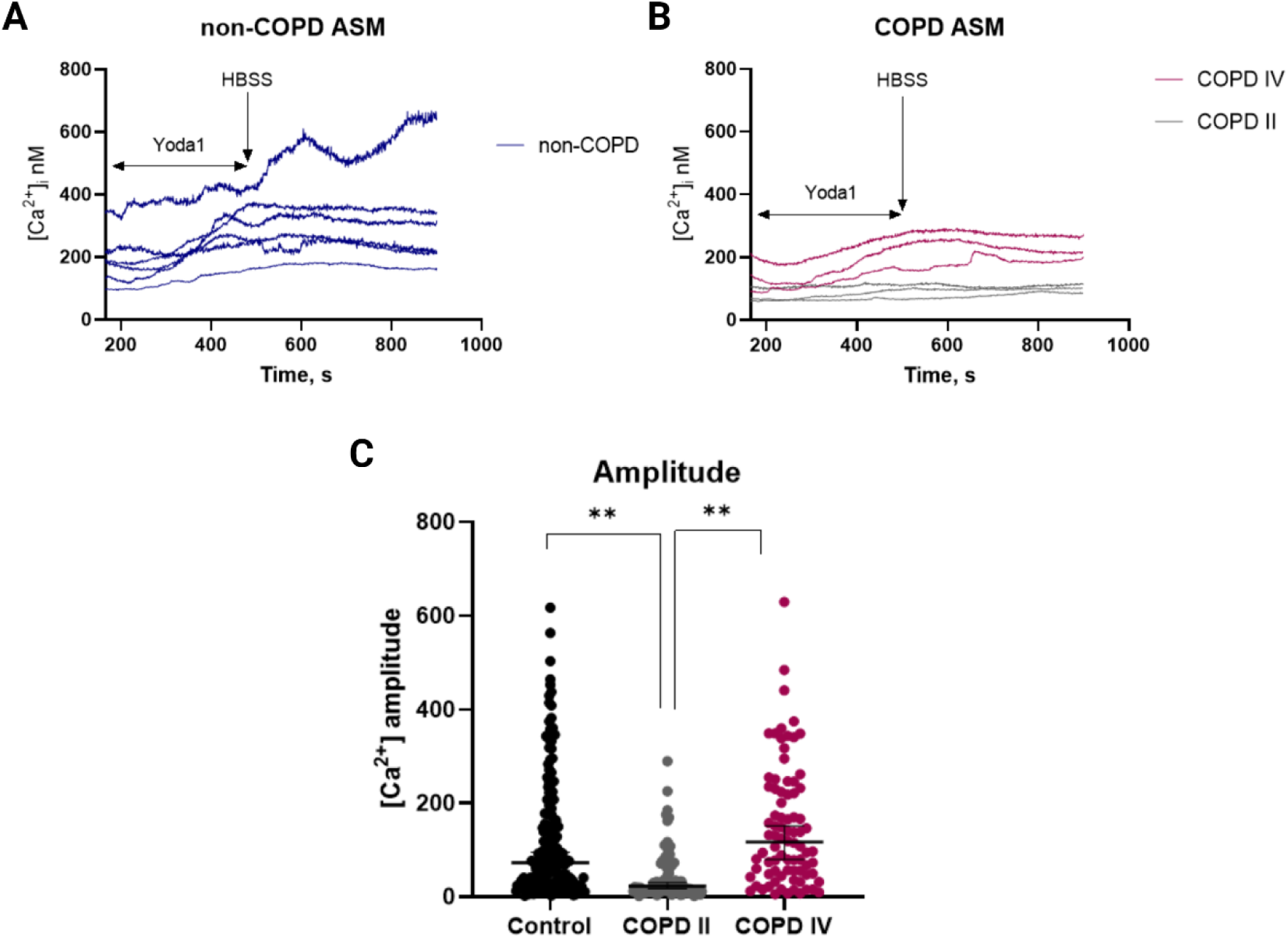
Effect of Yoda1 on intracellular Ca^2+^ concentration in non-COPD and COPD stage II and IV ASM. Intracellular Ca^2+^ concentration of ASM was measured over time using the Ca^2+^ indicator Fura-2/AM. Yoda1 was added at 165 seconds and maintained for 6 minutes before cells were washed with HBSS for 7 minutes; values are means of 24-32 cells per well plate from each donor. (A) intracellular Ca^2+^ concentration over time after stimulation of non-COPD ASM with Yoda1, (n=6); (B) intracellular Ca^2+^ concentration over time after stimulation of COPD II and COPD IV ASM with Yoda1, (n=3 per COPD subgroup); (C) the amplitude of cell response to Yoda1 stimulation measured as the difference between the maximum peak value and the value at the first moment of Yoda1 stimulation. Each dot is a value calculated for an individual ASM cell response (n=177, from 6 non-COPD patients; n=95, from 3 COPD II patients and n=82, from 3 COPD IV patients). Values are median with 95% confidence interval. Data were analyzed using a linear mixed-effects regression model to account for multiple measurements per donor; *p < 0.05. [Ca^2+^]i, intracellular Ca^2+^ concentration; ASM, airway smooth muscle; COPD, chronic obstructive pulmonary disease; HBSS, Hanks’ Balanced Salt solution.

### Yoda1 downregulates ECM gene and protein expression in ASM

We hypothesized that ECM-ASM crosstalk may be disrupted in COPD through aberrant Piezo channel function. Having investigated the impact of COPD and mechanical forces (stretch) on Piezo expression and functionality, we next examined the role of Piezo1 activation in regulating ECM production. We stimulated ASM with Yoda1 and measured gene expression of ECM and ECM-related genes (*LOX*, *LOXL1*, *LOXL2*, *Col1A*, *Col3*, *Fbln1*, *BMP1*, and *periostin*), involved in the fibrillogenesis of collagen and organization of ECM. We combined non-COPD and COPD ASM to test the effect of Yoda1 on ECM gene expression. *Col1A*, *Fbln1* and *periostin* were downregulated in ASM treated with Yoda1 compared to non-treated cells (Figure 7 A, B, C). There was no difference in *Col3A*, *BMP1, LOX*, *LOXL1* and *LOXL2* gene expression between ASM treated with vehicle vs. Yoda1 (Figure 7 D-H). The separate analysis of gene expression within non-COPD and COPD groups treated with Yoda1 did not reveal differences between the groups. However, we did observe that ASM from COPD II patients tended to respond more to Yoda1 stimulation compared to COPD IV and control ASM, however groups were too small for statistical analysis (Figure S2 A-F).

**Figure 7.**
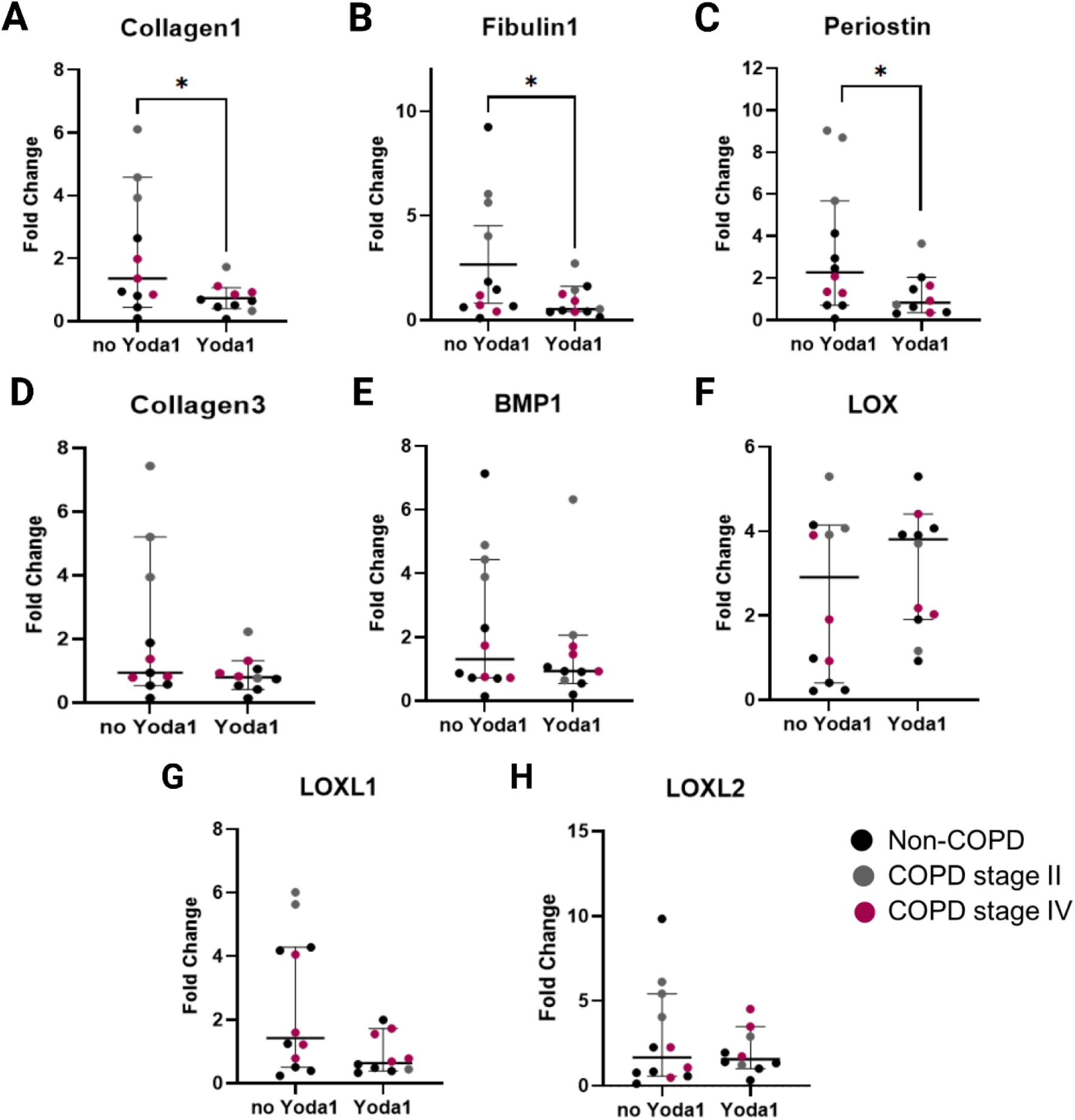
The effect of Yoda1 stimulation on ECM-related gene expression in ASM. ASM were stimulated with Yoda1 for 24 hours. Gene expression was analyzed using qPCR. Results are presented as relative expression to *16S*. (A) *Collagen1*; (B) *fibulin1*; (C) *periostin*; (D) *Collagen3*; (E) *BMP1*; (F) *LOX*; (G) *LOXL1*; (H) *LOXL2*. Data are expressed as median with 95% confidence interval, (n=12). The difference between ASM stimulated with and without Yoda1 was analyzed using the Wilcoxon test. *p < 0.05. Abbreviations: ASM, airway smooth muscle; BMP-1, bone morphogenetic protein; ECM, extracellular matrix; LOX, lysyl oxidase protein; LOXL1/2 – lysyl oxidase-like protein1/2.

In line with the observed changes on gene expression level, Collagen 1 and periostin protein expression was downregulated in ASM cells treated with Yoda1 compared to untreated cells (Figure 8 A, B). No differences were observed in Collagen 3 or BMP1 protein expression between treated and untreated ASM cells (Figure 8 C, D). Due to the small sample size, we could not perform separate statistical analyses for protein expression within the non-COPD and COPD groups treated with Yoda1. Nevertheless, we observed that ASM cells from all COPD patients exhibited lower periostin expression in response to Yoda1 stimulation compared to both the non-COPD and untreated groups (Figure S3 A-D).

**Figure 8.**
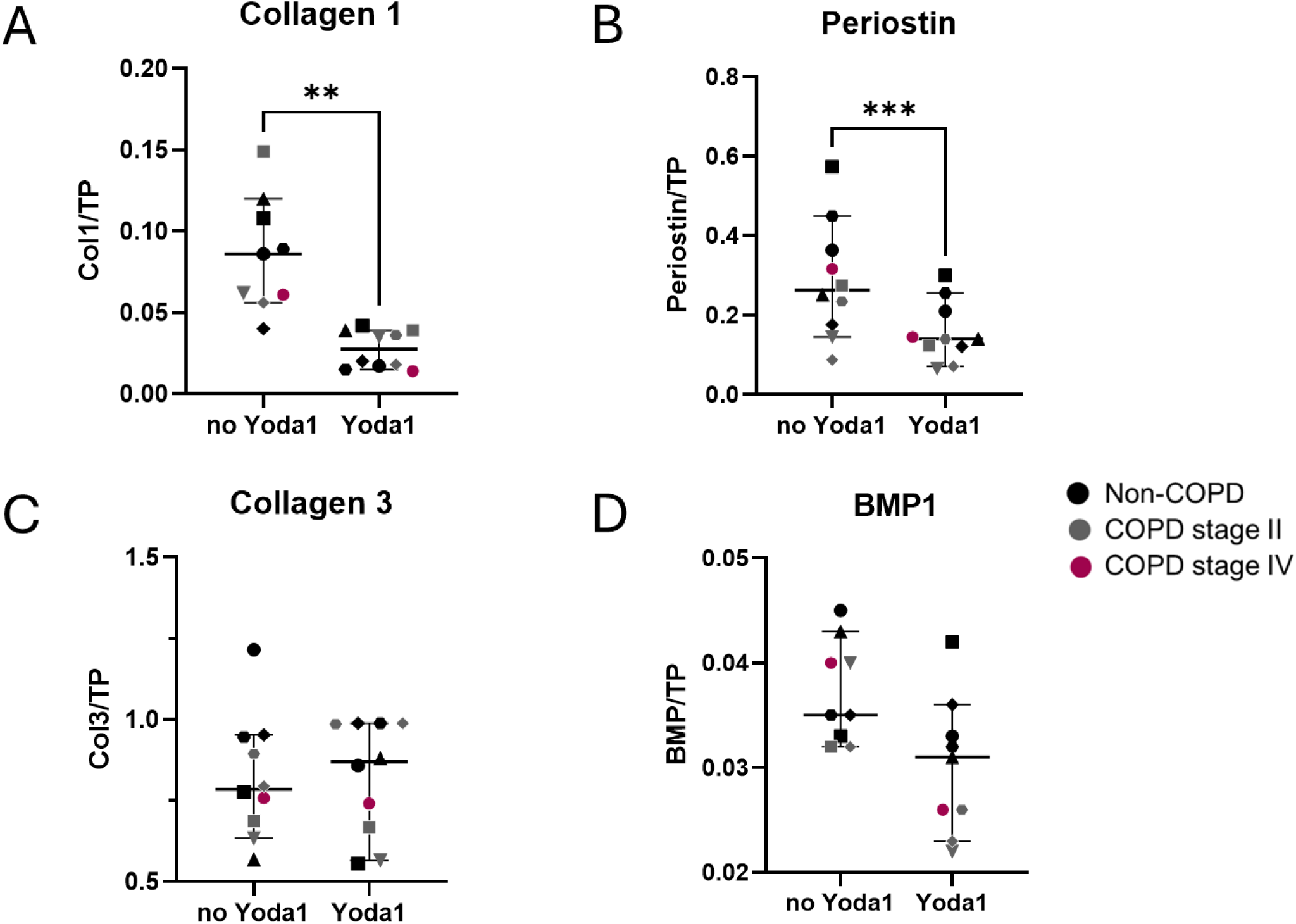
The effect of Yoda1 stimulation on ECM-related protein expression in ASM. ASM were stimulated with Yoda1 for 48 hours. Protein expression was analyzed using WB. Results are presented as relative expression to TP. (A) Collagen 1; (B) Periostin; (C) Collagen 3; (D) BMP1. Data is expressed as median with 95% confidence interval, (n=10). Each dot represents an individual donor, with the shape corresponding to a unique donor. The difference between ASM stimulated with and without Yoda1 was analyzed using the Wilcoxon or paired t-test, depending on the normality test. *p<0.05. Abbreviations: BMP1, bone morphogenetic protein: ECM, extracellular matrix: TP, total protein.

## Discussion

To our knowledge, this is the first study to examine the expression and activation of Piezo channels in ASM in the context of COPD and stretch. We report the presence and localization of both Piezo1 and Piezo2 in small airways, ASM bundles and in the epithelial layer and we observed less Piezo2 protein expression in the small airways, including the ASM bundles and epithelial layer in COPD IV compared to non-COPD lung tissue. When ASM cells were subjected to stretch *Piezo1* gene expression was upregulated in comparison to non-stretched ASM cells. This effect was more enhanced in non-COPD compared to COPD ASM cells. At the same time, Piezo2 protein expression was downregulated upon stretching but there was no difference between COPD and non-COPD. Upon Piezo activation with Yoda1, ASM cells from COPD II patients showed a significantly lower response in terms of intracellular Ca^2+^ influx compared to the COPD IV and non-COPD ASM cells. In addition, Yoda1 stimulation resulted in lower gene expression of *Col1A*, *Fbln1* and *periostin* and protein expression of collagen 1 and periostin compared to non-treated ASM cells. Altogether, our findings indicate there is altered expression and response to Piezo activation in COPD, which might be a consequence of altered mechanical stimuli such as disrupted alveolar attachments, stretch and ECM remodeling (Figure 9).

**Figure 9.**
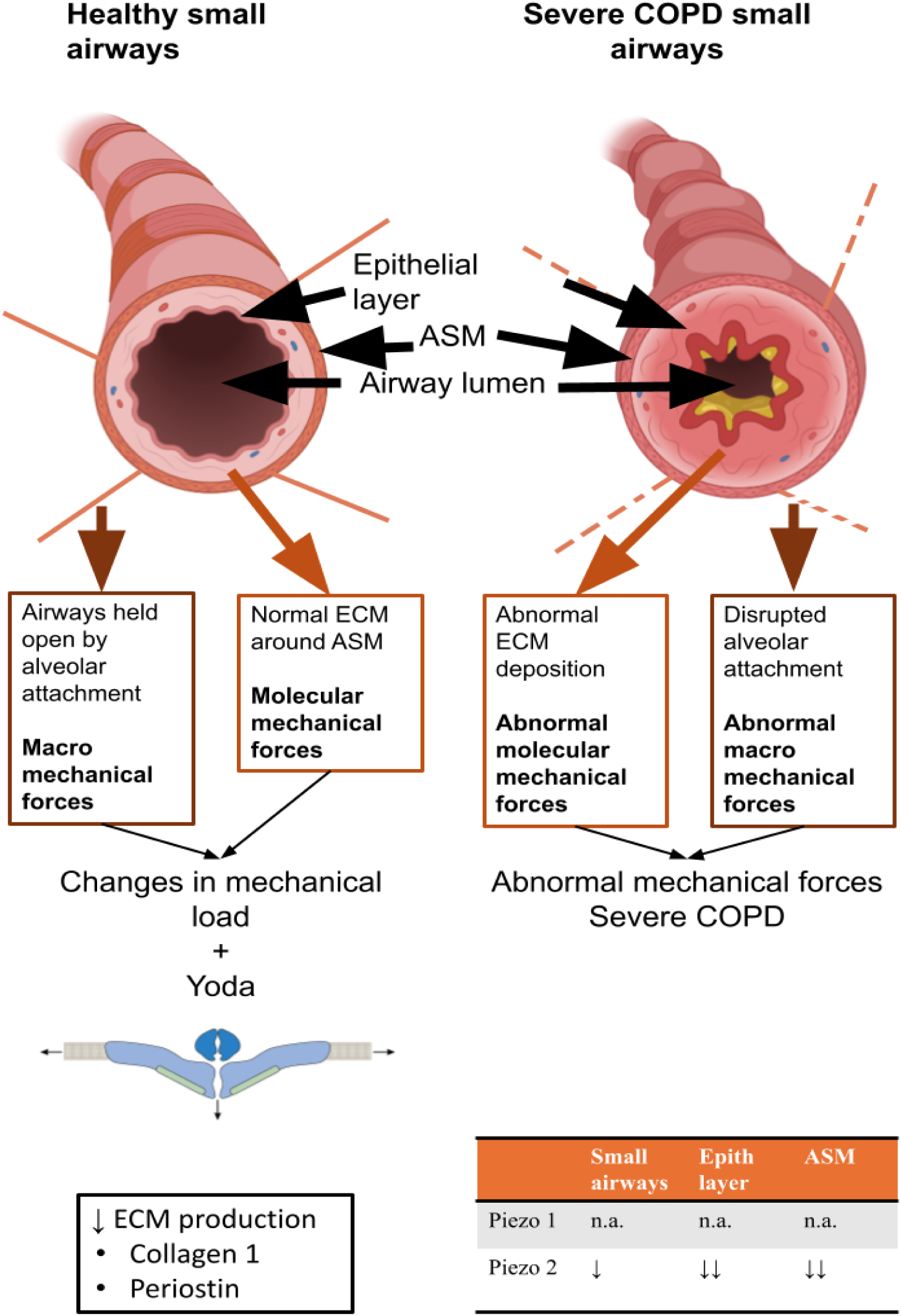
Schematic representation of the effects of COPD and mechanical forces on Piezo expression in small airways. In the small airways, increased mechanical load, both at the macroscopic level (due to alveolar attachments and airflow) and at the molecular level (due to ECM composition) can lead to Piezo activation. This, in turn, triggers a negative feedback loop, resulting in the downregulation of ECM production. In severe COPD, various factors including disruptions in alveolar attachments and abnormal ECM deposition alter mechanical forces, leading to changes in Piezo expression and activation within the small airways. Abbreviations: ASM, airway smooth muscle; ECM, extracellular matrix.

One of our main findings was lowered Piezo2 protein expression in COPD IV lung tissue. The downstream impact of these changes remains to be determined. However, downregulation of these channels in COPD may be compensated by promotion of Ca^2+^ influx in lung cells including ASM, and could change contraction and or relaxation and migration.^38^ A recent study has shown that activation of Piezo1 by Yoda1 in rat ASM resulted in decreased cell stiffness and traction force.^26^ We hypothesize that changes in Ca²⁺-regulated pathways involving Piezo channels may contribute to airway remodeling in COPD. Conversely, abnormal mechanical forces in already remodeled COPD airways may alter Piezo channel activation and expression. *In vivo*, the lungs are continuously exposed to mechanical forces such as shear stress, compression, and stretch during breathing and pulmonary circulation.^39^ ASM sense various physical forces and convert them to intracellular signals that underlie normal physiology of the respiratory system which may also contribute to the pathogenesis of COPD.^38^ Interestingly, we found that stretching affects gene regulation after 24 h and protein production at 48 h for Piezo channels in ASM. In our model, the parameters set for the stretching cycles were designed to mimic normal breathing patterns in healthy lungs. In COPD cells, we did not find a difference in Piezo1 expression upon stretch. Although this could be due to limited power, this could also indicate that the stretching forces that apply to the ASM *in vivo* in the lungs of patients with COPD may have imprinted these cells to then result in resistance to mechanical stimuli. In COPD ASM localizes within fibrotic tissue in the small airways where there are elevated ECM levels.^40^ Consequently, the augmented mechanical forces may induce cellular adaptation, enhancing the capacity of ASM to endure heightened mechanical stresses. Therefore, COPD ASM could become less sensitive to the normal physiological stretching forces mimicked in our experimental setup. After 24 h, stretching did not change *Piezo2* gene expression. However, after 48 h Piezo2 protein was downregulated in stretched ASM cells compared to non-stretched, to a similar extent in the COPD and non-COPD ASM cells. The difference in gene and protein expression of Piezos could be a result of timing of sampling in our experiment and the complexity of transcription and translation regulation.

Piezo protein expression patterns were slightly different between ASM cells in culture and ASM bundles in lung tissue *in vivo* from non-COPD and COPD patients. Non-stretched COPD ASM did not differ in both gene and protein expression of Piezo1 and Piezo2 compared to non-COPD cells. Piezo1 expression percentage area and intensity also were not different in ASM layers from COPD lungs, while Piezo2 intensity in ASM layers from COPD IV patients was significantly lower compared to non-diseased tissue. These discrepancies between isolated cells and intact tissue may reflect the effect of cellular/extracellular environment, where switching from a soft tissue environment to the greater stiffness of the *in vitro* cell culture plates may alter Piezo expression. Given that Piezos are biomechanical channels, differences in stiffness likely has a vast impact on Piezo expression and activation.^39^ This aspect remains to be further explored in the context of airway cells.

Along with expression patterns of Piezo in COPD ASM, we investigated functional differences of Piezo in COPD. ASM derived from COPD II patients showed a lower response to Yoda1 in terms of [Ca^2+^]_i_, compared to COPD IV and non-COPD ASM. In isolated ASM cells expression of Piezo1 gene and protein in COPD cells was similar to non-COPD and no obvious differences were present between COPD II and COPD IV cells. This discrepancy between Ca^2+^ response and Piezo1 expression may reflect Piezo1 channel sensitivity and/or the impact of the extracellular environment on Piezo1 activation: factors that remain to be further explored. Of note, in the ASM bundles in the lung tissue sections, a trend towards lower area and intensity of staining for Piezo1 was detected in the COPD IV group compared to non-COPD. The somewhat lower expression of Piezo1 in COPD IV ASM bundles *in vivo* and active Ca^2+^ influx *in vitro* may result from a compensatory mechanism to maintain Ca^2+^ influx levels in COPD IV ASM via this channel as seen *in vitro*. The dissimilarity observed in the Ca^2+^ response between COPD stage II and IV may be attributable to variances in the expression of Piezo channels on the ASM cells at the different stages. Additionally, this dissimilarity might be an indication that in stage II COPD the ASM cells are still mechanoresponsive and attempting to respond to the aberrant mechanical environment in the tissue, however by stage IV the ASM cells response to the mechanical environment has been exhausted and the expression of Piezo is subsequently abrogated. Alternatively, in stage II COPD, the tissue may contain a mix of both functional and dysfunctional ASM cells, while by stage IV, only the more resilient, functional cells may survive. As a result, the remaining ASM cells in stage IV may exhibit Ca²⁺ influx levels that are more active than in stage II and relatively comparable to those of healthy cells.

We propose that Piezos play a role in ECM remodeling in COPD, given that Ca^2+^ channels are known to be associated with regulation of ECM production.^41^ Intracellular Ca^2+^ is a second messenger, regulating a number of cellular functions. For example, in fibrosis and tumor metastasis, dysregulated matrix remodeling is associated with another mechanosensitive Ca^2+^-permeable channel, transient receptor potential vanilloid type 4, and disruptions of Ca^2+^ homeostasis.^41^ We observed lower expression of *Collagen1, fibulin1* and *periostin* genes and collagen 1 and periostin proteins in ASM treated with Yoda1 compared to untreated ASM. A possible explanation is that the reduced ECM expression results from negative feedback of increased extracellular forces present in the remodeled or fibrotic areas in the COPD lung that activate Piezo1 in an attempt to reduce ECM production. As with Ca^2+^ influx in COPD II ASM after Yoda1 stimulation, ASM from COPD II also tended to have a more pronounced difference in ECM gene expression in response to Yoda1 stimulation compared to the control and COPD IV ASM. However, we acknowledge that our study was not sufficiently powered for a proper statistical comparison between different COPD severities and thus this needs further investigation.

Although we have reported several novel findings about Piezo1 and Piezo2 in human lung tissue, there are limitations. An important challenge was that due to the absence of a commercially available agonist for Piezo2, investigation into the functional role of Piezo2 was not possible without siRNA or other protein knockdown approaches that complicate the model. Furthermore, cells were cultured on two-dimensional stiff tissue culture well plates, which do not reflect the mechanical environment of the lung. This may limit the interpretation of the results and *in vitro* to *in vivo* translation of our findings, as cell behavior and Piezo channel functionality likely differs in a 3-dimensional environment.

In conclusion, this study is the first to explore the role of Piezo channels in relation to stretch forces and the presence of COPD in ASM, covering changes in the gene expression and protein levels as well as functional studies. These findings highlight the potential role of Piezo channels in healthy and COPD lung physiology. Further advances will be needed to fully understand the contribution of Piezo channels to the underlying mechanisms driving airway remodeling and disease pathogenesis of COPD.

## Abbreviation List

ASM: Airway Smooth Muscle
BMP1: Bone Morphogenetic Protein 1
Col1A: Collagen 1A
Col3: Collagen 3
COPD: Chronic Obstructive Pulmonary Disease
ECM: Extracellular Matrix
Fbln1: Fibulin 1
FBS: Fetal Bovine Serum
FEV1: Forced Expiratory Volume in 1 second
FVC: Forced Vital Capacity
GOLD: Global Initiative for Chronic Obstructive Lung Disease
HBSS: Hanks’ Balanced Salt Solution
IHC: immunohistochemistry
LOX: Lysyl Oxidase
LOXL1: Lysyl Oxidase-like 1
LOXL2: Lysyl Oxidase-like 2
PBS: Phosphate Buffered Saline
RT: Room Temperature
SD: Standard Deviation
TP: Total Protein
TRP: Transient Receptor Potential (channels)
TRPV4: Transient Receptor Potential Vanilloid type 4

