## Supplemental materials for "Mechanosensitive Piezo Channels Contribute to Airway Changes in Chronic Obstructive Pulmonary Disease"

### Supplementary figures

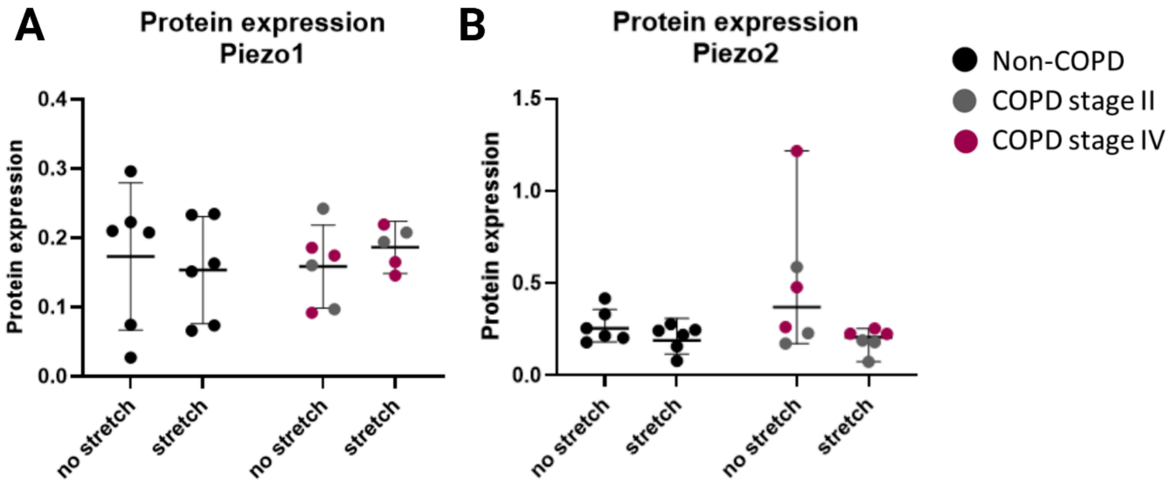


**Figure S1.** ASM derived from COPD patients did not have altered Piezo protein expression in response to stretch. ASM derived from non-COPD and COPD patients were stretched in a cell system FlexCell for 48 hours. The protein expression of Piezo was analyzed using WES system. (A) Piezo1 protein expression in non-stretched and stretched ASM from non-COPD and COPD patients (n=6); (B) Piezo2 protein expression in non-stretched and stretched ASM from non-COPD and COPD patients. Data are expressed as median with 95% (n=6). The difference between non-stretched and stretched groups was analyzed using the Wilcoxon matched-pairs test, and the difference between non-COPD and COPD ASM was analyzed using the Mann-Whitney U test. *$p < 0.05$. Abbreviations: ASM, airway smooth muscle; COPD, chronic obstructive pulmonary disease.

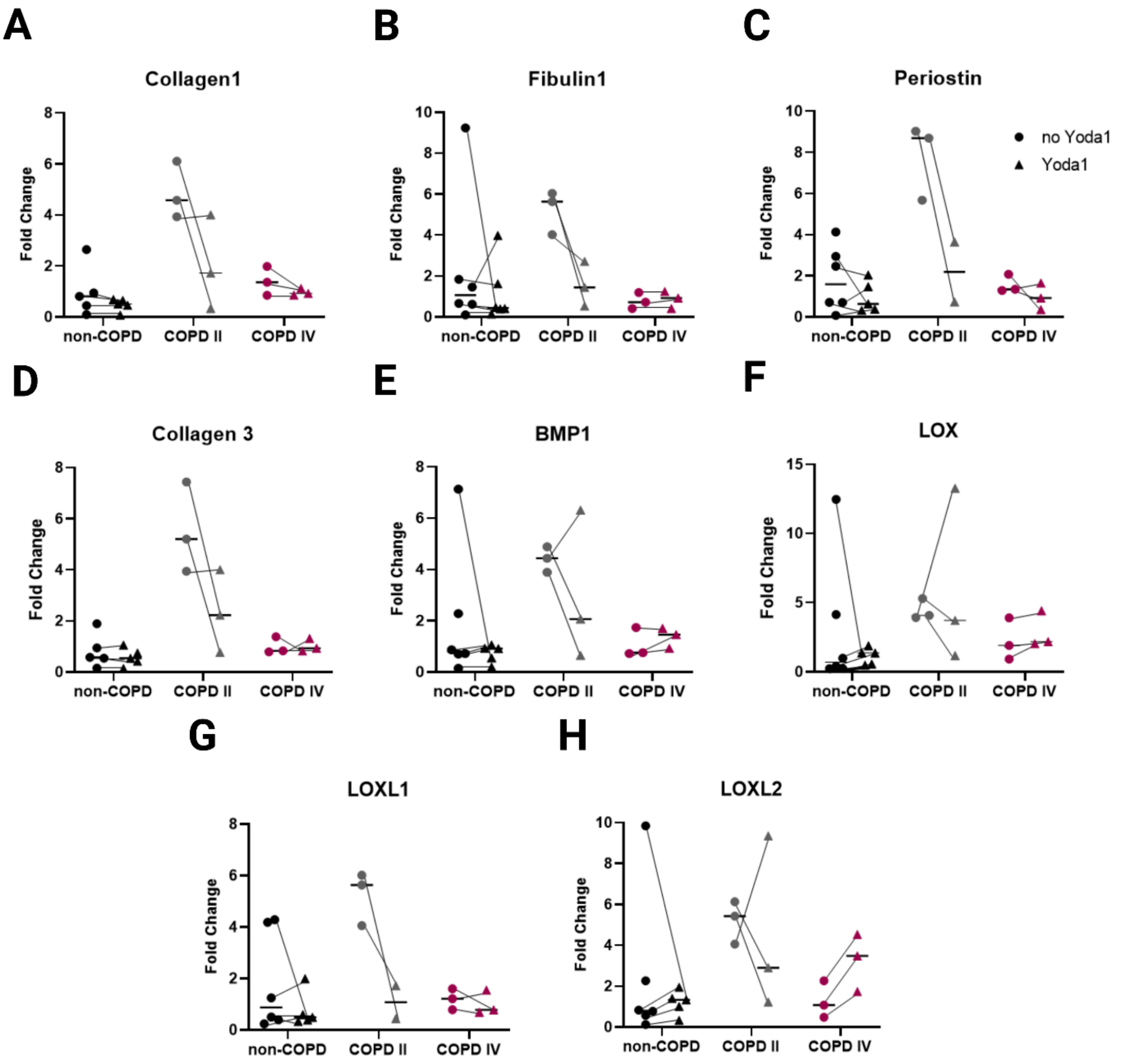


**Figure S2.** The effect of Yoda1 stimulation on ECM-related gene expression in non-COPD, COPD II and COPD IV ASM separately. ASM were stimulated with Yoda1 for 24 hours. Gene expression was analyzed using qPCR. (A) *Collagen1*; (B) *fibulin1*; (C) *periostin*; (D) *Collagen 3*; (E) *BMP1*; (F) *LOX*; (G) *LOXL1*; (H) *LOXL2*. Data are expressed as median, (n=6 for control and n=3 & 3 for COPD II/COPD IV). Abbreviations: BMP-1, bone morphogenetic protein; COPD, chronic obstructive pulmonary disease; ECM, extracellular matrix; LOX, lysyl oxidase protein; LOXL1/2 – lysyl oxidase-like protein1/2.

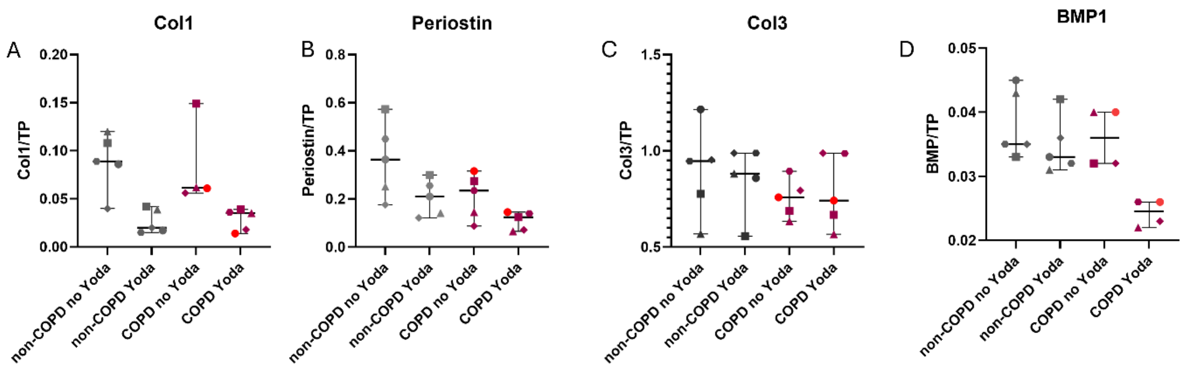


**Figure S3.** The effect of Yoda1 stimulation on ECM-related protein expression in non-COPD and COPD ASM. ASM were stimulated with Yoda1 for 24 hours. Protein expression was analyzed using WB. Results are presented as relative expression to TP. (A) Collagen 1; (B) Collagen 3; (C) Periostin; (D) BMP1. Data are expressed as median with 95% confidence interval, (n=5). Abbreviations: BMP1, bone morphogenetic protein: ECM, extracellular matrix: TP, total protein.
